# Differential Biofilm Susceptibility and Potent Isavuconazole Post-Antifungal Effect Distinguish *Cutaneotrichosporon dermatis* from *Trichosporon asahii*

**DOI:** 10.64898/2026.08.30.748177

**Authors:** Tatsuya Yoshinouchi, Tomofumi Nakamura, Daisuke Mori, Jun-ichirou Yasunaga, Yasuhito Tanaka

## Abstract

*Cutaneotrichosporon dermatis* (formerly *Trichosporon dermatis*) is a basidiomycetous yeast-like fungus known to cause summer-type hypersensitivity pneumonitis, although its virulence in humans remains poorly understood. We performed morphological and molecular identification of an isolate from the sputum and blood cultures of an immunocompromised patient, together with pathogenicity assessment using a *Galleria mellonella* model, biofilm formation/eradication assays, antifungal susceptibility testing, drug combination effects, and the post-antifungal effect (PAFE), compared with *Trichosporon asahii*. The isolate was identified as *C. dermatis* by ITS/IGS1 sequencing, supported by phylogenetic analysis. Growth of *C. dermatis* increased more at 37□ than at 25□. In the *Galleria mellonella* assay, *C. dermatis*, *T. asahii*, and *Candida albicans* each showed dose-dependent pathogenicity at sufficiently high inocula, although *Rhizopus oryzae* was the most potent pathogen on a per-CFU basis. *C. dermatis* formed biofilms that were more completely inhibited by terbinafine (TRB) and amphotericin B (AmB) than azole agents, which showed only partial inhibitory activity even at high concentrations. Susceptibility testing showed relatively strong susceptibility to AmB and azole agents. In the TRB and azole combination assay, the fractional inhibitory concentration index (FICI) was below 0.5, indicating synergy. Isavuconazole (ISC) showed a markedly stronger PAFE than the other azole agents tested. These findings indicate that although azoles show only partial efficacy against its biofilm, *C. dermatis* can still cause invasive infection, and that azole monotherapy or TRB and azole combination therapy, aided by the potent PAFE of ISC, may represent effective treatment options.

## Introduction

The family Trichosporonaceae, which includes the genus *Trichosporon*, has undergone taxonomic reorganization with advances in molecular phylogenetic analysis, and several clinically important species have been transferred to the genera *Cutaneotrichosporon* and *Apiotrichum* [1]. *Cutaneotrichosporon dermatis* was originally classified within the *Cryptococcus humicola* complex, but in 2001 it was established as a new species within the genus *Trichosporon* together with *Trichosporon debeurmannianum* [2], and was subsequently transferred to the genus *Cutaneotrichosporon* following the integrated phylogenetic classification [1].

Among *Trichosporon* species, *T. asahii* is well known as the major causative agent of disseminated trichosporonosis and has been reported to cause fatal fungemia in immunocompromised patients such as those with hematologic malignancies or neutropenia [3]. In contrast, reports of *C. dermatis* (formerly *T. dermatis*) as a cause of human infection are limited, although sporadic cases of invasive infection in immunocompromised patients have been reported, including fungemia in an infant with fever of unknown origin [4] and fungemia after chemotherapy in a patient with Burkitt lymphoma [5]. In a multicenter study, only a small proportion of *Trichosporon* isolates recovered from blood cultures were identified as *C. dermatis*, and its clinical significance remains incompletely established [6]. In the present study, the organism was likewise isolated from the sputum and blood cultures of a patient who became immunocompromised following high-dose corticosteroid therapy administered for an underlying disease.

Additionally, *C. dermatis* is classified as one of the major antigenic serotypes (serotype I) of summer-type hypersensitivity pneumonitis (SHP), a form of hypersensitivity pneumonitis that occurs predominantly in summer in Japan [7, 8]. Serotypes are determined serologically by cell slide agglutination and ELISA using type-specific antisera raised against surface polysaccharide antigens [7]. In serotyping analyses of *Trichosporon* isolates recovered from the homes of SHP patients, 97.7% of serotype I isolates were identified as *C. dermatis* [9], suggesting that this species may be important not only as a focus of invasive infection but also as an antigen involved in allergic airway sensitization. Therefore, when this fungus is detected in the sputum of an immunocompromised patient, it is clinically important to consider whether this represents simple airway colonization or a pathological process (progression from colonization to invasion, or induction of an allergic reaction).

Regarding pathogenicity, experimental evaluation of a clinical isolate of *C. dermatis* in a murine model has already been reported [10], but pathogenicity assessment using the insect infection model *Galleria mellonella*, as employed in the present study, has not previously been performed for this species. *The Galleria mellonella* model has already been established for pathogenicity assessment and antifungal efficacy testing in other *Trichosporon* species such as *T. asahii*, *T. asteroides*, and *T. inkin* [11], but it has not yet been applied to *C. dermatis*.

Antifungal susceptibility data for *C. dermatis* also remain limited to environmental isolates such as those from coral reefs [12], and systematic susceptibility profiles using multi-drug panels in human clinical isolates are scarce. In *T. asahii*, biofilm formation has been reported to contribute to drug resistance [13, 14, 15], and the usefulness of biofilm-targeted combination therapy (evaluated by checkerboard-based FICI assessment) has been demonstrated [16]; however, no combination-therapy data are currently available for *C. dermatis*.

We therefore performed a comprehensive characterization of a *C. dermatis* isolate obtained from the sputum and blood cultures of an immunocompromised patient, including morphological and molecular identification (including phylogenetic analysis), pathogenicity assessment using the *Galleria mellonella* infection model, antifungal susceptibility testing, and evaluation of drug combination effects, and compared these findings with those of *T. asahii* to clarify the clinical and microbiological characteristics of this species.

## Materials and Methods

### 1. Fungal isolates, reference strains, and media

Yeast-like fungi were isolated from the sputum culture and blood culture of an immunocompromised patient. *C. dermatis* isolated from the blood culture was used in this study. As reference strains, *C. dermatis* (NBRC-102675), *T. asahii* (NBRC-10844), *Candida albicans* (NBRC-1594), and *Rhizopus oryzae* (ATCC-56965) were used. Sabouraud dextrose agar (SDA) was prepared using meat peptone (5 g/L; Nacalai Tesque, Kyoto, Japan), casein peptone (5 g/L; Nacalai Tesque), glucose (40 g/L; Nacalai Tesque), and agar (1.5% w/v; Nacalai Tesque). After autoclaving, chloramphenicol (Cp; FUJIFILM Wako, Osaka, Japan) and kanamycin (K; FUJIFILM Wako) were added. Separately, RPMI 1640 medium (Nissui Pharmaceutical, Tokyo, Japan) was prepared without NaHCO_3_ and adjusted to pH 7.0 using 0.165 M MOPS (3-morpholinopropane-1-sulfonic acid; Nacalai Tesque) and NaOH (Nacalai Tesque). This MOPS-buffered RPMI medium was sterilized by filtration through a 0.22 µm filter and supplemented with chloramphenicol (Cp) and kanamycin (K).

### 2. Species identification and phylogenetic analysis

DNA extraction was performed as previously described [17]. Briefly, fungal material cultured on SDA was harvested, rapidly frozen in liquid nitrogen, and disrupted using a BioMasher II (Nippi, Tokyo, Japan). Total DNA was extracted from the resulting homogenate using TRIzol reagent (Invitrogen, Thermo Fisher Scientific) according to the manufacturer’s instructions and stored at −80□. For species identification, the following gene regions were amplified by PCR using 10–50 ng of template DNA: for the internal transcribed spacer (ITS) region, primers NS7F (5′-GAG GCA ATA ACA GGT CTG TGA TGC-3′) and ITS4R (5′-TCC TCC GCT TAT TGA TAT GC-3′); for the IGS region, primers 26SF (5′-ATC CTT TGC AGA CGA CTT GA-3′) and 5SR (5′-AGC TTG ACT TCG CAG ATC GG-3′). PCR was performed using KOD One PCR Master Mix (TOYOBO, Osaka, Japan) in a 40 µL reaction, using an Eppendorf Mastercycler thermal cycler (Eppendorf, Hamburg, Germany). Cycling conditions were: initial denaturation at 98□ for 2 min (1 cycle), followed by 35 cycles of denaturation at 98□ for 10 s, annealing at 55□ for 5 s, and extension at 68□ for 15 s, with a final extension at 68□ for 1 min. Species identification was confirmed by comparison of the resulting ITS and IGS1 sequences with MycoBank reference sequences on the basis of Score, Probability, and Similarity values, using a similarity threshold of ≥99%.

For phylogenetic analysis, reference sequences obtained from GenBank (National Center for Biotechnology Information) were used, prioritizing sequences derived from ex-type strains wherever possible. The ITS dataset comprised 32 sequences including the present isolate (18 Cutaneotrichosporon species, 11 Trichosporon species, and two Apiotrichum species used as an outgroup: A. montevideense and A. mycotoxinovorans). The IGS1 dataset comprised 18 sequences including the present isolate (5 Cutaneotrichosporon species, 11 Trichosporon species, and A. mycotoxinovorans as the sole outgroup). All GenBank accession numbers are listed in Tables S1 (ITS) and S2 (IGS1). Sequences for each region were aligned separately using MUSCLE [18] as implemented in MEGA version 12.1.2 [19]. The best-fit nucleotide substitution model was selected on the basis of the Bayesian Information Criterion (BIC) using the "Find Best DNA Model (ML)" function in MEGA, and the Tamura–Nei model was selected for both datasets. Phylogenetic trees were constructed by the maximum likelihood method, and topological robustness was assessed by bootstrap analysis (1,000 replicates), with trees rooted using the outgroups described above.

### 3. Morphological observation by light and scanning electron microscopy

For light microscopy, the growth morphology of *C. dermatis* and *T. asahii* cultured in MOPS-buffered medium was observed using a BZ-X810 microscope (Keyence, Japan). For scanning electron microscopy (SEM), isolates cultured on SDA plates were fixed in 2% glutaraldehyde in pH 7.0 phosphate buffer for 48 h at room temperature with shaking, dehydrated through a graded ethanol series (50%, 75%, 90%, 95%, and 100%; 15 min per step at room temperature with shaking), dried by the t-butyl alcohol sublimation method, and coated with platinum. Samples were observed using a JSM-IT300LA scanning electron microscope (JEOL, Tokyo, Japan). To visualize fungal adhesion to host cells, A549 cells (1.5×10^5^ cells/mL) were seeded onto 18×18 mm coverslips in chamber slide II dishes (IWAKI, Japan) containing DMEM supplemented with 10% FBS, penicillin, and kanamycin, and incubated overnight at 35□ in 5% CO□. *C. dermatis* or *T. asahii* was then added to a final concentration of 1.0×10^5^ CFU/mL, and the medium was replaced with DMEM containing 1% FBS, chloramphenicol, and kanamycin; cultures were incubated for a further 24 h at 35□ in 5% CO_2_. Samples were fixed with 2% glutaraldehyde in 0.1 M phosphate buffer (pH 7.4) for 48 h and processed for SEM as described above.

### 4. Growth kinetics assay

Isolates of *C. dermatis*, *T. asahii*, and *C. albicans* were inoculated into 96-well plates in MOPS-buffered RPMI medium and incubated at 25□ and 37□. OD_530_ was measured at 0, 12, 24, 36, 48, 60, and 72 h to assess the change in fungal biomass over time. Medium-only wells served as the negative control. All absorbance readings were performed with a FLUOstar Omega plate reader (BMG Labtech, Germany).

### 5. Pathogenicity assessment using the *Galleria mellonella* infection model

The species identity of the purchased *Galleria mellonella* larvae (Amapure Farm, Chiba, Japan) was confirmed in advance by gene sequence analysis. Larvae were frozen in liquid nitrogen, minced, and homogenized using a tissue homogenizer. Total DNA was extracted using TRIzol reagent (Invitrogen) according to the manufacturer’s instructions. A partial region of the mitochondrial cytochrome c oxidase subunit I (COI) gene was amplified using the universal primers LCO1490 (5′-GGT CAA CAA ATC ATA AAG ATA TTG G-3′) and HCO2198 (5′-TAA ACT TCA GGG TGA CCA AAA AAT CA-3′) [20]. Cycling conditions were: initial denaturation at 94□ for 3 min, followed by 35 cycles of denaturation at 94□ for 30 s, annealing at 48 for 45 s, and extension at 72□ for 1 min, with a final extension at 72□ for 5 min. The amplification product was sequenced and subjected to a BLAST search against GenBank, showing 100% identity to *Galleria mellonella* (Accession No. OY292312.1). The Galleria mellonella infection model was performed as previously described [11, 21]. Larvae weighing 200–300 mg were selected (PBS control, n = 10; each inoculated group, n = 17–18). Larvae were inoculated with PBS (control), *C. dermatis* (5×10^4^, 5×10^5^, and 1×10^6^ CFU/larva), *T. asahii* (5×10^4^, 5×10^5^, and 1×10^6^ CFU/larva), *C. albicans* (5×10^4^, 5×10^5^, and 1×10^6^ CFU/larva), or *R. oryzae* (1×10^3^, 5×10^3^, and 1×10^4^ CFU/larva) via the left proleg using a microsyringe (Hamilton 701N, 10 µL), and observed at 35°C for 120 h. Survival was calculated by the Kaplan–Meier method using EZR [22], and between-group comparisons were made using the log-rank test.

### 6. Antifungal susceptibility testing

The minimum inhibitory concentration (MIC) of each antifungal agent was determined by broth microdilution in accordance with Clinical and Laboratory Standards Institute (CLSI) document M27, 4th edition (2017), following a previously described method [23]. Two-fold serial dilutions of each antifungal agent were prepared in MOPS-buffered RPMI 1640 medium. For the fungal inoculum, blastoconidia of *C. dermatis* and arthroconidia of *T. asahii* were harvested from cultures grown on SDA for 2–5 days and suspended in PBS containing 0.5% Tween 80 (Nacalai Tesque, Japan). The suspension was counted using a hemocytometer, added to 96-well flat-bottom plates, and adjusted to a final inoculum concentration of 1.0×10^3^ CFU/mL for yeasts. Plates were incubated at 35□, and the MIC was determined visually after 24 h (extended to 48 h for isolates with insufficient growth at 24 h). The MIC was defined as the lowest drug concentration (mg/L) inhibiting visible growth. The 50% and 90% inhibitory concentrations (IC50, IC90) were determined by WST-1 staining (Dojindo Laboratories, Kumamoto, Japan) and absorbance at 440 nm, using a FLUOstar Omega plate reader, compared with positive (no drug) and negative (medium only) controls. Voriconazole (VRC), itraconazole (ITC), posaconazole (PSC), and terbinafine (TRB) were purchased from Tokyo Chemical Industry (Tokyo, Japan); amphotericin B (AmB) and isavuconazole (ISC) from Selleck Chemicals (Houston, TX, USA); micafungin (MCF) from Cayman Chemical (Ann Arbor, MI, USA). Stock solutions (5–20 mM in DMSO or another appropriate solvent) were stored at −80 and diluted to working concentration in MOPS-buffered RPMI immediately before use.

### 7. Evaluation of drug combination effects

The drug combination assay was performed based on a previously described method [17]. Briefly, combinations of TRB and each azole agent at the concentrations to be tested were prepared in flat-bottom 96-well plates using MOPS-RPMI medium. Cells of *C. dermatis* were inoculated at 5.0 × 10^3^ CFU/mL and incubated at 35 for 48 h. The MIC (mg/L) was determined by visual observation of *C. dermatis* growth. IC_50_ and IC_90_ (mg/L) were determined by quantifying the viability of *C. dermatis* at OD440 nm after WST-1 staining. The percentage growth inhibition by the drug combination was calculated by dividing each value by that of the positive control (*C. dermatis* in drug-free MOPS-RPMI). The fractional inhibitory concentration index (FICI) for each combination tested was calculated according to a previously described method [24], and for each of the MIC, IC_50_ (WST-1), and IC_90_ (WST-1) datasets, FICI ≤ 0.5 was defined as synergy, 0.5 < FICI ≤ 1 as additive, 1 < FICI ≤ 4 as indifferent, and FICI > 4 as antagonistic [16].

### 8. Biofilm formation, inhibition, and eradication assays

Biofilm inhibition was determined using the crystal violet (CV) staining assay [17, 25]. Cells of *C. dermatis* 1 and *T. asahii* were seeded at 1.0×10^6^ CFU/mL in flat-bottomed collagen I-coated 96-well plates together with the tested drugs and incubated at 35 for 24 h. The wells were washed twice with 100 μL of PBS and stained with 100 μL of 0.1% CV solution for 30 min at room temperature (RT). The wells were then washed three times with 200 μL of PBS and air-dried at RT. Bound CV was extracted by adding 100 μL of 30% acetic acid and incubating with shaking for 15 min at RT. An 80-μL aliquot of the resulting solution was transferred to a fresh 96-well plate, and absorbance was measured at 620 nm (OD_620_). Biofilm inhibition by each drug was expressed as the relative ratio of OD_620_ compared with the positive control (fungal growth in drug-free MOPS-RPMI) and the negative control (MOPS-RPMI medium only, without fungal inoculation). Biofilm eradication was assessed as previously described [17, 25]. Briefly, cells of *C. dermatis* 1 and *T. asahii* were seeded at 5.0×10^5^ CFU/mL in 100 μL of drug-free MOPS-RPMI in 96-well plates and incubated at 35 for 24 h to allow biofilm formation. The tested drugs were then added at each concentration, and the plates were incubated for a further 24 h at 35□. The eradicating activity of each drug was determined using the CV staining assay described above.

### 9. Post-antifungal effect (PAFE) assay

The post-antifungal effect (PAFE) of each antifungal agent against *C. dermatis* and *T. asahii* was determined using a drug-exposure/washout method [17, 26]. Briefly, fungal suspensions (3.0×10^5^ CFU/mL) were exposed to each antifungal agent (VRC, ISC, ITC, PSC, or TRB at 0.5–32 mg/L; AmB at 0.13–16 mg/L) for either 6 h or 24 h at 35 with shaking at 200 rpm, as two independent exposure conditions. After exposure, a 10-µL aliquot of the fungal suspension was washed twice with 1000 µL of PBS to remove residual drug, resuspended in 300 µL of drug-free MOPS-buffered RPMI medium, and divided among three wells of a 96-well plate. Fungal regrowth was monitored 24 h after washout by measuring cell viability (WST-1 assay, OD_440_), and results are expressed as the ratio relative to a drug-free control.

### 10. Statistical analysis

For the PAFE assay, the drug-exposed group (6 h or 24 h) was compared with the drug-free control group for each antifungal agent using Student’s t-test. Survival curves in the *Galleria mellonella* model were compared using the Kaplan–Meier method and the log-rank test. All statistical analyses were performed with EZR (Jichi Medical University, Tochigi, Japan) [22]. *P* values < 0.05 was considered statistically significant.

## Results

### 1. Species identification and molecular phylogenetic placement

We attempted identification by matrix-assisted laser desorption/ionization time-of-flight mass spectrometry (MALDI-TOF MS; Vitek MS), but this method could not distinguish *C. dermatis* from *C. mucoides*, consistent with previous reports of MALDI-TOF misidentification between these closely related species [28, 29, 30]. We therefore performed molecular analysis instead. The results of sequencing the rDNA ITS and IGS1 regions of the isolate are shown in Table S4. The isolate was identified as *Cutaneotrichosporon dermatis* with 100% identity in the ITS region (query coverage 79.9%) and 100% identity in the IGS1 region (coverage 100%). Phylogenetic trees based on the ITS and IGS1 regions, constructed using GenBank reference sequences, are shown in Fig. 1. In both trees, the present isolate formed a monophyletic clade with reference sequences, supporting its identification as *C. dermatis* at the molecular phylogenetic level, independent of the BLAST/MycoBank similarity search results. Notably, a clear difference in resolution was observed between the two markers (ITS and IGS1). In the ITS tree, the bootstrap value at the node closest to the isolate was 63%, decreasing further to 45% and 43% at deeper nodes, indicating relatively low support for branching among closely related sequences. In contrast, in the IGS1 tree, the terminal clade containing the isolate had a bootstrap value of 100%, resolving relationships within and among closely related species with high resolution. This finding is consistent with previous reports indicating that, within the genera *Trichosporon*/*Cutaneotrichosporon*, the ITS region alone has limited discriminatory power among closely related species, whereas the IGS1 region provides higher resolution [27]. In both trees, sequences diverging on long basal branches, separate from the main clade containing the isolate, were observed; this likely reflects the reorganization and separation of the genus *Cutaneotrichosporon* from *Trichosporon* following the integrated phylogenetic classification by Liu *et al*. [1].

**Fig. 1.**
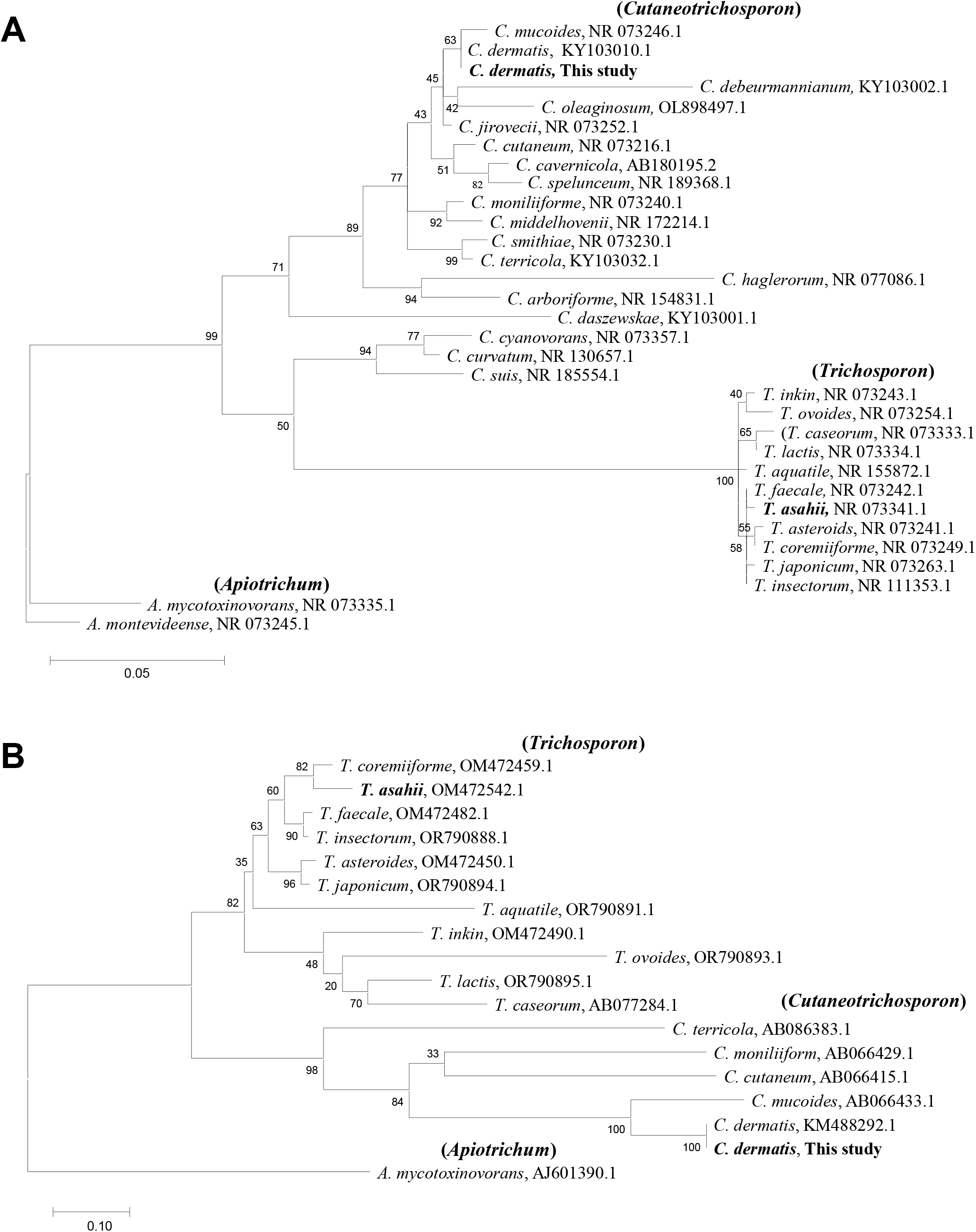
Phylogenetic trees based on the rDNA ITS and IGS1 region. Phylogenetic trees based on the rDNA ITS region (**A**) and IGS1 region (**B**), constructed by the maximum likelihood method following multiple alignment with MUSCLE in MEGA12. The present isolate (bold) formed a monophyletic clade with GenBank reference sequences in both trees. Numbers at each node indicate bootstrap values (%) from 1,000 replicates. Genus-level groupings (*Cutaneotrichosporon*, *Trichosporon*, *Apiotrichum*) are indicated. Accession numbers of the reference sequences used are listed in Tables S1 (ITS) and S2 (IGS1).

### 2. Morphological findings

The morphological findings presented in Fig. 2 showed a clear contrast between *C. dermatis* (clinical isolate) and *T. asahii* (NBRC-10844). On SDA, *C. dermatis* formed smooth, glossy, milky-white yeast-like colonies, whereas *T. asahii* formed characteristic cerebriform colonies with a wrinkled, uneven surface and a dry appearance reflecting filamentous growth. In MOPS-RPMI culture, *C. dermatis* showed predominantly clustered, budding yeast-like oval-to-round blastoconidia, with wavy hyphal structures under phase-contrast microscopy. In contrast, *T. asahii* showed a dense network of true and pseudohyphae composed of elongated filamentous structures occupying the entire field, with relatively few yeast-like blastoconidia. A similar contrast was observed by scanning electron microscopy (SEM). *C. dermatis* exhibited spherical-to-oval cells aggregated in a cauliflower-like structure, whereas *T. asahii* formed a dense fibrous network of hyphae extending in multiple directions, with some images suggestive of segmented arthroconidia formation. This morphological contrast is consistent with the taxonomic background of these organisms. The genus *Trichosporon* is characterized by the formation of true and pseudohyphae and the production of arthroconidia through their segmentation, whereas some closely related genera, including *Cutaneotrichosporon*, show relatively yeast-like growth (predominantly blastoconidia); this has been cited as one of the phenotypic bases for the generic reorganization in the 2015 integrated phylogenetic classification [1]. The original description of *C. dermatis* similarly reported that the species shows predominantly yeast-like growth composed of blastoconidia, with hyphal formation being sparse [2]. The morphological findings obtained in the present study were in agreement with these previous descriptions.

**Fig. 2.**
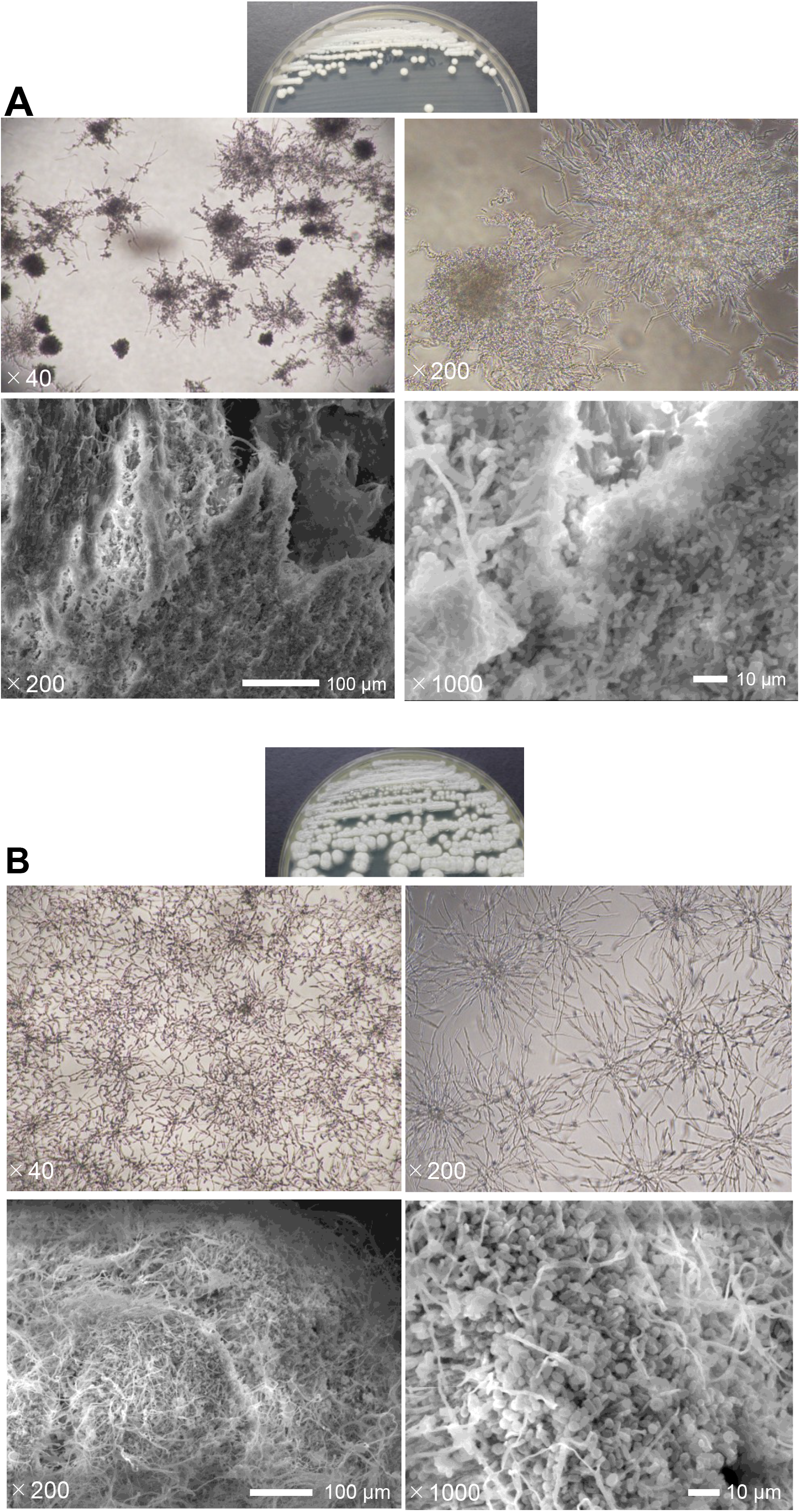
Morphological findings of *C. dermatis* and *T. asahii*. Morphological findings of *C. dermatis* (**A**) and *T. asahii* (**B**) are shown from top to bottom: colony morphology on Sabouraud dextrose agar (SDA); phase-contrast microscopy of cultures in MOPS-RPMI medium; and scanning electron microscopy (SEM), at the magnifications indicated. Scale bars and magnifications are shown in each image.

### 3. Growth kinetics over time

We examined the growth kinetics of *C. dermatis*, *T. asahii*, and *C. albicans* at 25□ and 37□ in MOPS-RPMI buffered medium by measuring OD_530_ over time (Fig. 3). *C. dermatis* showed a temperature-dependent growth pattern, with a greater increase in OD_530_ over time and more rapid biomass accumulation at 37□ than at 25□ (Fig. 3A); microscopic examination at this stage showed clustered, yeast-like cells and wavy hyphal structures consistent with the morphology described in Fig. 2. In contrast, *T. asahii* showed similar growth patterns at 25□ and 37□ (Fig. 3B), with a dense hyphal network again consistent with Fig. 2. *C. albicans*, used as a growth control, showed a more rapid and pronounced increase in OD_530_ at both temperatures than either *C. dermatis* or *T. asahii* (Fig. 3C). When compared directly at each temperature, *C. dermatis* and *T. asahii* showed comparable growth at 25□ (Fig. 3D), whereas at 37°C *C. dermatis* reached a substantially higher OD_530_ than *T. asahii* from 24 h onward (Fig. 3E).

**Fig. 3.**
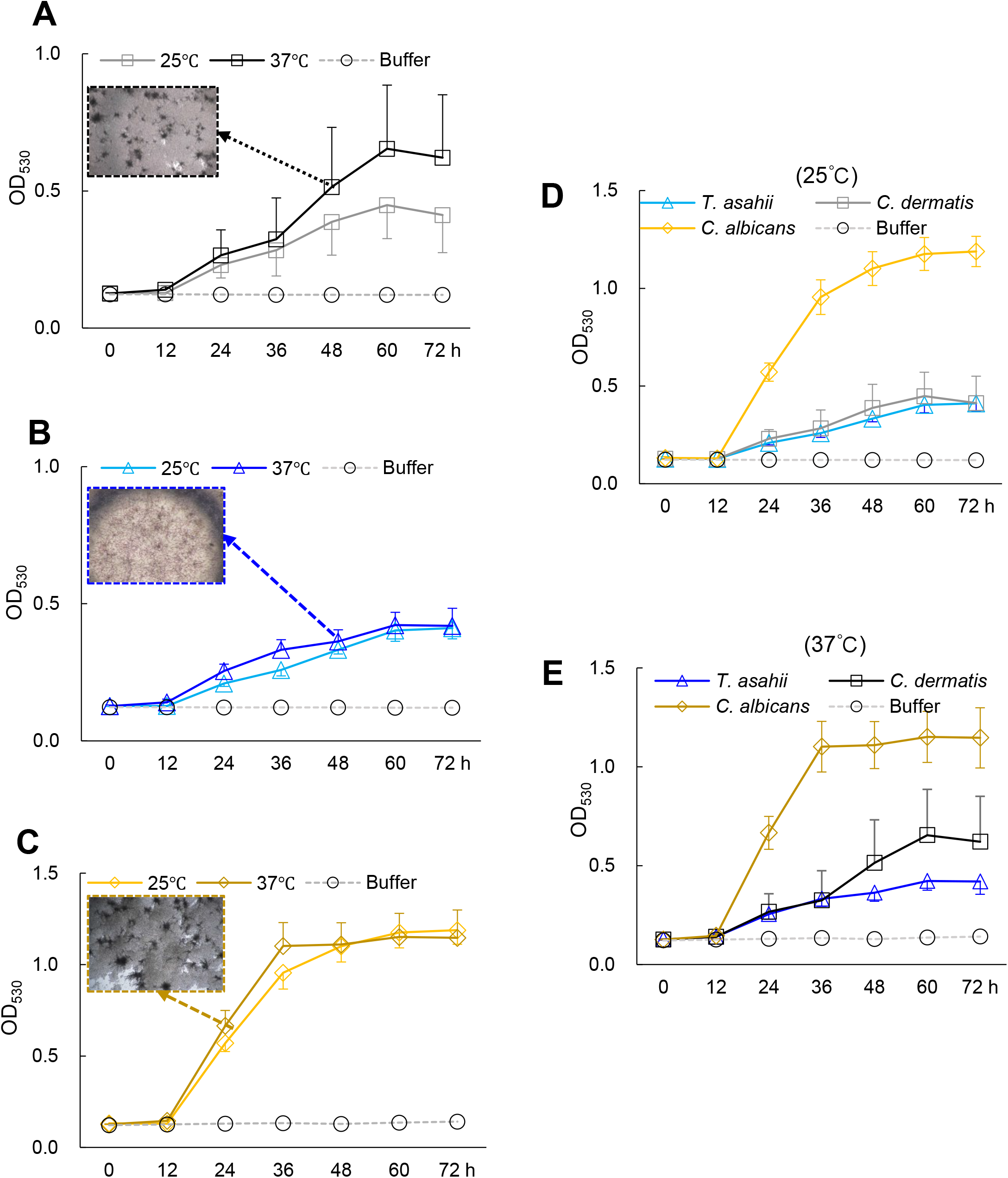
Growth kinetics of *C. dermatis*, *T. asahii*, and *C. albicans* over time in MOPS-RPMI buffered medium. Growth kinetics were assessed by measuring OD_530_ at 25□ and 37□ every 12 h for up to 72 h. (**A**) Growth kinetics of *C. dermatis* at 25□ (gray line) and 37□ (black line), with a representative image of the culture at 48 h. (**B**) Growth kinetics of *T. asahii* at 25□ (light blue line) and 37□ (blue line), with a representative image at 48 h. (**C**) Growth kinetics of *C. albicans* at 25□ (yellow line) and 37□ (dark yellow line), with a representative image at 24 h. (**D**, **E**) Comparison of *T. asahii* and *C. dermatis* growth at 25□ (**D**) and 37□ (**E**). Gray dashed lines indicate the medium-only negative control. Data shown are average values of two independent experiments.

### 4. Antifungal susceptibility and drug combination effects of *C. dermatis*

The antifungal susceptibility profiles of *C. dermatis* 1 (clinical isolate), *C. dermatis* 2 (NBRC-102675), and *T. asahii* are shown in Table 1. All three isolates showed relatively low MIC values and potent susceptibility to amphotericin B (AmB) [31], voriconazole (VRC) [32], itraconazole (ITC) [33], posaconazole (PSC) [34], and isavuconazole (ISC) [35]. In contrast, susceptibility to terbinafine (TRB) [36] was intermediate, with higher MIC values than AmB and the azole agents, and both species showed markedly elevated MIC values to micafungin (MCF) (>16 mg/L), suggestive of intrinsic resistance. Among the three isolates, susceptibility of *C. dermatis* 1 was comparable to that of *T. asahii*, whereas the susceptibility of *C. dermatis* 2 to the azole agents was stronger than that of *C. dermatis* 1 and *T. asahii*.

**Table 1.** The susceptibility of *C. dermatis* and *T. asahii* to the antifungals. MIC was defined as the lowest drug concentration (mg/L) that caused complete visual inhibition of fungal growth. IC_50_ and IC_90_ values represent the drug concentrations (mg/L) that inhibited 50% and 90% of fungal growth, respectively, as determined by cell viability OD_440_ following WST-1 staining. Data shown are representative of three independent experiments. Abbreviations are as follows: VRC, voriconazole; ITC, itraconazole; PSC, posaconazole; ISC, isavuconazole; MCF, micafungin; TRB, terbinafine; AmB, amphotericin B.

| Species | Assay endpoint | Drugs |  |  |  |  |  |  |
| --- | --- | --- | --- | --- | --- | --- | --- | --- |
|  |  | VRC | ISC | ITC | PSC | AmB | TRB | MCF |
| *C. dermatitis* 1 clinical isolate | IC<sub>50</sub> (OD<sub>530</sub>) | 0.031 | 0.063 | 0.016 | 0.25 | 0.13 | 0.5 | >16 |
|  | IC<sub>90</sub> (OD<sub>530</sub>) | 0.25 | 0.13 | 0.063 | 0.5 | 0.25 | 1 | >16 |
|  | IC<sub>50</sub> (WST-1) | 0.25 | 0.13 | 0.016 | 0.25 | 0.13 | 1 | >16 |
|  | IC<sub>90</sub> (WST-1) | 0.5 | 0.5 | 0.031 | 0.5 | 0.25 | 2 | >16 |
|  | MIC | 0.13 | 0.5 | 0.25 | 0.5 | 0.13 | 1 | >16 |
| *C. dermatitis* 2 reference strain | IC<sub>50</sub> (OD<sub>530</sub>) | 0.031 | 0.016 | 0.016 | 0.063 | 0.063 | 0.25 | 16 |
|  | IC<sub>90</sub> (OD<sub>530</sub>) | 0.063 | 0.13 | 0.031 | 0.13 | 0.13 | 1 | >16 |
|  | IC<sub>50</sub> (WST-1) | 0.016 | 0.016 | 0.016 | 0.031 | 0.063 | 0.5 | 16 |
|  | IC<sub>90</sub> (WST-1) | 0.031 | 0.063 | 0.016 | 0.063 | 0.13 | 2 | >16 |
|  | MIC | 0.031 | 0.031 | 0.016 | 0.063 | 0.13 | 1 | >16 |
| *T. asahii* | IC<sub>50</sub> (OD<sub>530</sub>) | 0.063 | 0.13 | 0.063 | 0.13 | 0.063 | 1 | >16 |
|  | IC<sub>90</sub> (OD<sub>530</sub>) | 0.063 | 0.25 | 0.13 | 0.25 | 0.5 | 2 | >16 |
|  | IC<sub>50</sub> (WST-1) | 0.063 | 0.25 | 0.063 | 0.25 | 0.063 | 1 | >16 |
|  | IC<sub>90</sub> (WST-1) | 0.063 | 0.5 | 0.13 | 0.5 | 0.25 | 4 | >16 |
|  | MIC | 0.13 | 0.5 | 0.13 | 0.25 | 0.25 | 4 | >16 |

Additionally, the results of the combination assay of TRB with each azole agent against *C. dermatis* 1 are shown in Table 2. For the TRB + VRC combination, synergy (FICI ≤0.5) was observed at all endpoints: the 50% inhibitory concentration (IC_50_, FICI 0.31), the 90% inhibitory concentration (IC_90_, FICI 0.37), and the visual endpoint (MIC, FICI 0.31). For the TRB + ISC combination, stronger synergy was also observed at all endpoints: IC_50_ (FICI 0.38), IC_90_ (FICI 0.37), and MIC (FICI 0.28). For the TRB + ITC combination, equally strong synergy was observed at all endpoints: IC_50_ (FICI 0.28), IC_90_ (FICI 0.28), and MIC (FICI 0.31). For the TRB + PSC combination, synergy was shown at IC_50_ (FICI 0.32), IC_90_ (FICI 0.38), and MIC (FICI 0.38). For the TRB + AmB combination, an additive (0.5 < FICI ≤ 1) or indifferent (1 < FICI ≤ 4) interaction was shown at IC_50_ (FICI 1.0), IC_90_ (FICI 1.0), and MIC (FICI 1.5). These results suggest that combination therapy with TRB and an azole agent may be a useful treatment option for *C. dermatis* infection.

**Table 2.** *In vitro* combination effects of TRB with azoles and AmB against *C. dermatis*.

|  | WST-1 (OD<sub>440</sub>) |  | MIC |
| --- | --- | --- | --- |
|  | IC<sub>50</sub> | IC<sub>90</sub> |  |
| TRB | 0.5 | 0.5 | 0.5 |
| VRC | 0.0078 | 0.031 | 0.016 |
| FIC index | 0.31 | 0.37 | 0.31 |
| TRB | 0.13 | 0.25 | 0.13 |
| ISC | 0.0078 | 0.016 | 0.0078 |
| FIC index | 0.38 | 0.37 | 0.28 |
| TRB | 0.5 | 1 | 0.5 |
| ITC | 0.0019 | 0.0019 | 0.016 |
| FIC index | 0.28 | 0.28 | 0.31 |
| TRB | 0.13 | 0.25 | 0.13 |
| PSC | 0.016 | 0.063 | 0.063 |
| FIC index | 0.32 | 0.38 | 0.38 |
| TRB | 0.13 | 0.25 | 0.5 |
| AmB | 0.13 | 0.25 | 0.13 |

| FIC index | 1.0 | 1.0 | 1.5 |
| --- | --- | --- | --- |

Combination effects of TRB with VRC, ISC, ITC, or PSC, and TRB with AmB against *C. dermatitis* by the checkerboard method. The 50% and 90% inhibitory concentrations (IC<sub>50</sub>, IC<sub>90</sub>; determined by WST-1 assay), the visually determined MIC (mg/L), and the FICI are shown for each combination. FICI was calculated as described in Materials and Methods. Data shown are representative of two independent experiments.

### 5. Anti-biofilm formation and eradication of anti-fungal agents against *C. dermatis* and *T. asahii*

To examine biofilm formation by *C. dermatis* and *T. asahii*, fungal cells cultured on SDA or on A549 cell monolayers were observed by SEM, and the inhibitory and eradicative effects of antifungal agents on biofilm formation were examined using a CV assay (Fig. 4). On SDA, *C. dermatis* formed a filamentous structure composed of agglutinated hyphae and oval conidia, and *T. asahii* formed a mixture of hyphae and oval conidia without tight cohesion (Fig. 4A, D). In co-culture with A549 cells, however, *C. dermatis* hyphae penetrated into the cell monolayer, and mature conidia appeared fused with the cells, indicating invasive adhesion, whereas *T. asahii* showed weak adhesion, with only sparse contact points between the hyphae and the cells (Fig. 4B, E).

**Fig. 4.**
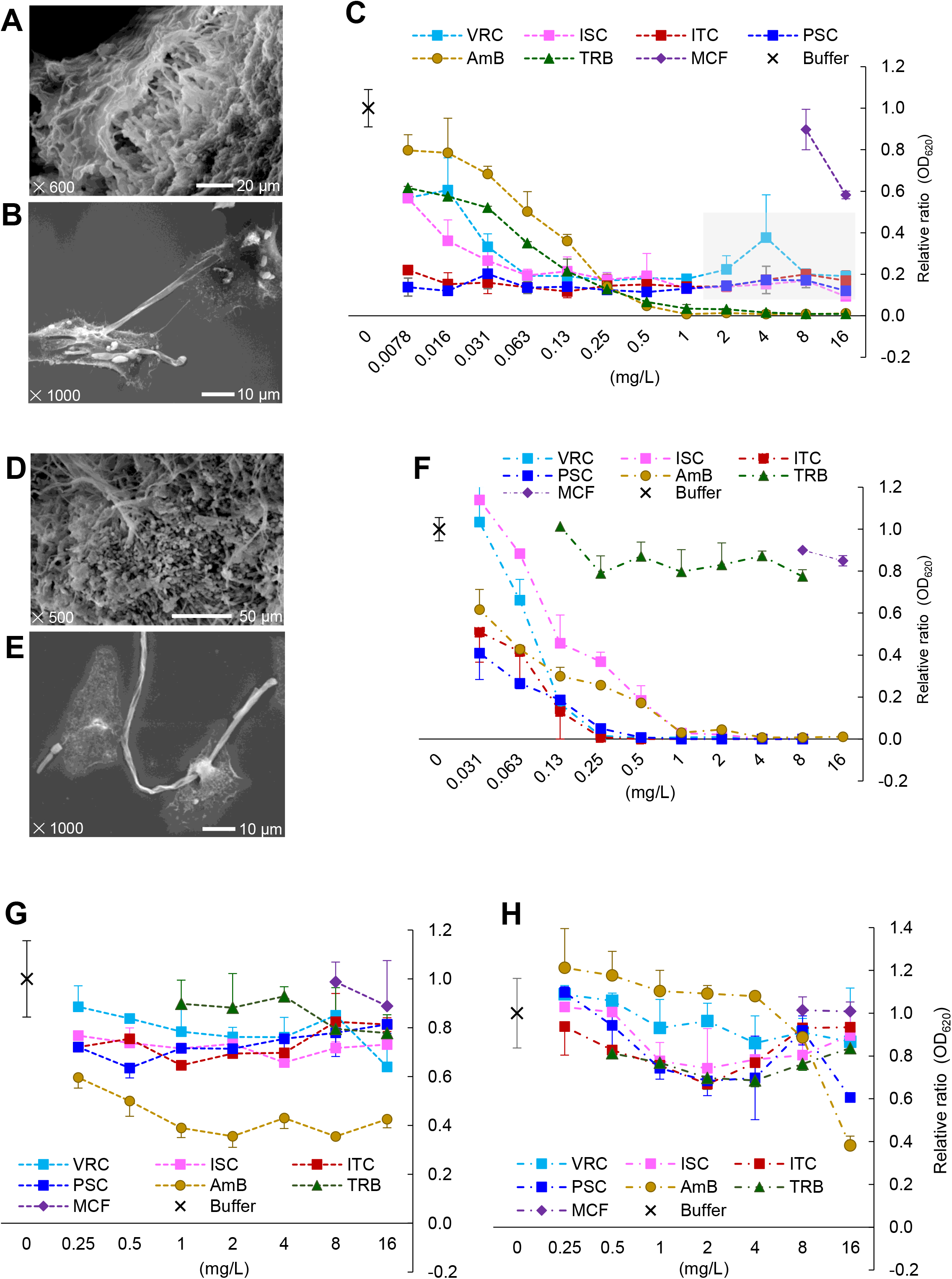
Biofilm formation, inhibition, and eradication by *C. dermatis* and *T. asahii*. (**A**, **B**) SEM of *C. dermatis* cultured on SDA (**A**) and on A549 cell monolayers for 24 h (**B**). (**D**, **E**) Corresponding SEM images for *T. asahii* on SDA (**D**) and on A549 monolayers (**E**). (**C**, **F**) Relative biofilm biomass (OD_620_, crystal violet assay) for *C. dermatis* (**C**) and *T. asahii* (**F**) after treatment with VRC, ISC, ITC, PSC, AmB, TRB, or MCF during biofilm formation (inhibition assay). (**G**, **H**) Relative biofilm biomass for *C. dermatis* (**G**) and *T. asahii* (**H**) after treatment of a pre-formed, mature biofilm with each drug (eradication assay). Values in **C**, **F**, **G**, and **H** are expressed relative to the drug-free control (=1.0). All assays were performed in triplicate, and error bars indicate ± SD from at least two independent experiments. AmB, amphotericin B; ISC, isavuconazole; ITC, itraconazole; MCF, micafungin; PSC, posaconazole; TRB, terbinafine; VRC, voriconazole.

We next performed a CV assay to evaluate the effects of the tested antifungal agents on biofilm formation by *C. dermatis* and *T. asahii* (Fig. 4C, F). Interestingly, higher concentrations (>2 mg/L) of TRB and AmB achieved near-complete inhibition of biofilm formation by *C. dermatis*, whereas azole agents (ISC, ITC, and PSC) showed only partial inhibition, with OD620 plateauing at approximately 0.1–0.2 relative to the drug-free control across the 2–16 mg/L range rather than declining further; VRC showed similar partial inhibition at low-to-intermediate concentrations but then a marked, non-monotonic increase, rising to approximately 0.9 at 8 mg/L before declining again at 16 mg/L, resembling the paradoxical growth effect (Eagle effect) previously reported for antifungal agents [37, 38] (Fig. 4C). In contrast, MCF showed no inhibitory effect on biofilm formation by *T. asahii*, whereas TRB showed almost no inhibitory effect, and azole agents and AmB suppressed biofilm formation by *T. asahii*, in a dose-dependent manner, with complete inhibition at higher concentrations of azole agents (>1 mg/L) and AmB (>4 mg/L), unlike the pattern observed for *C. dermatis* (Fig. 4F). These results demonstrated that biofilm formation, and its inhibition by the tested antifungal agents, differed between *C. dermatis* and *T. asahii*.

We further evaluated the eradicating activity of the tested antifungal agents against pre-formed biofilms of *C. dermatis* and *T. asahii* (Fig. 4G, H). For *C. dermatis*, AmB reduced biofilm biomass by up to approximately 60% across the 0.25–16 mg/L concentration range, whereas the azole agents reduced OD_620_ by only around 20%, without a clear dose-dependent pattern; TRB showed a similarly modest effect, reducing biofilm biomass by up to approximately 20%, and MCF showed almost no eradicating activity (Fig. 4G). For *T. asahii*, AmB and PSC reduced biofilm biomass by approximately 60% and 40%, respectively, at 16 mg/L; TRB unexpectedly reduced biofilm biomass by approximately 20%, despite showing almost no inhibitory effect on biofilm formation itself, whereas MCF again showed no reduction in biofilm biomass (Fig. 4H). Overall, these results suggest that complete eradication of established biofilms is difficult to achieve for either species, and that the eradicating activity of a given drug does not necessarily correlate with its ability to inhibit initial biofilm formation.

### 6. Post antifungal effects of antifungal agents against *C. dermatis* and *T. asahii*

The post-antifungal effects (PAFE) of the tested antifungal agents against *C. dermatis* and *T. asahii*, reflecting their residual growth-inhibitory potency after drug removal, are shown in Fig. 5. Against *C. dermatis*, PSC suppressed fungal growth at high concentrations, and ISC produced the most pronounced and consistent suppression of regrowth after 6-h drug exposure, with significantly lower relative growth ratios than VRC, ITC, TRB, and AmB at 24 h post-washout, across all concentrations tested (Fig. 5A).

**Fig. 5.**
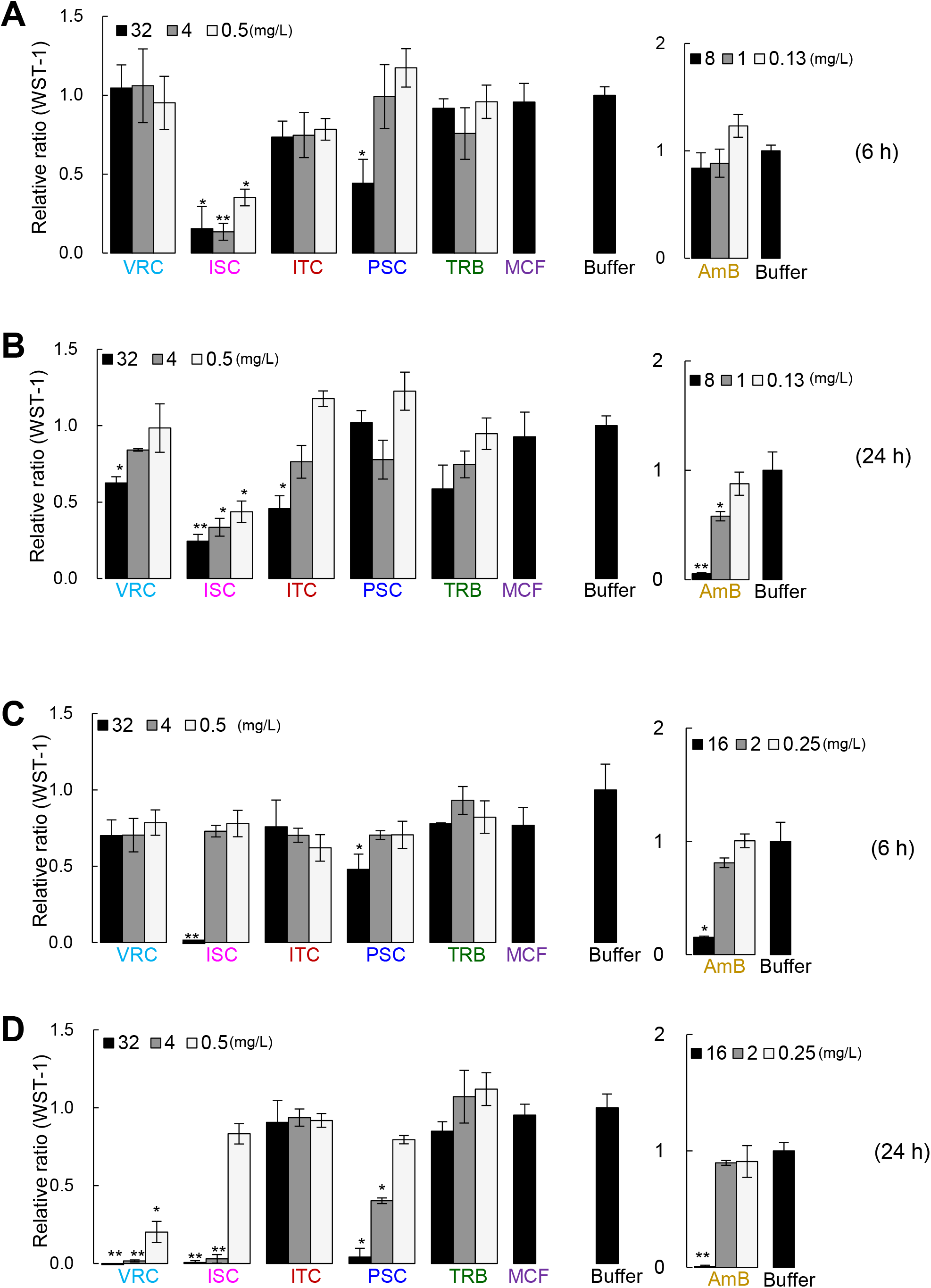
Post-antifungal effect (PAFE) of antifungal agents against *C. dermatis* and *T. asahii*. Fungal cells were exposed to each antifungal agent (VRC, ISC, ITC, PSC, TRB, AmB, and MCF) at 0.13–32 mg/L concentration for 6 h (**A**, **C**) or 24 h (**B**, **D**), washed, and monitored for regrowth 24 h after washout by WST-1 assay. Values are expressed as the ratio relative to a drug-free control (=1.0); lower values indicate stronger suppression of regrowth (a more potent PAFE). All assays were performed in triplicate, and error bars indicate ± SD from at least two independent experiments. Statistical significance was determined by Student’s *t*-test (*, *P* < 0.05 **, *P* < 0.005). AmB, amphotericin B; ISC, isavuconazole; ITC, itraconazole; MCF, micafungin; PSC, posaconazole; TRB, terbinafine; VRC, voriconazole.

After 24-h drug exposure, 32 mg/L VRC and ITC, 8 and 32 mg/L AmB, and all concentrations of ISC showed significant suppression of regrowth (Fig. 5B). Against *T. asahii*, VRC, ISC, and PSC showed pronounced suppression of regrowth at higher concentrations (4 and 32 mg/L), particularly after 24-h drug exposure, whereas ITC and TRB showed comparatively weaker post-antifungal effects (Fig. 5C, D). AmB showed a concentration-dependent PAFE against both species, with stronger suppression of regrowth at the highest concentration tested (8 mg/L for *C. dermatis* and 16 mg/L for *T. asahii*) (Fig. 5A–D).

These results indicate that ISC may exert prolonged residual growth-inhibitory activity against *C. dermatis* even after only 6 h of exposure at low concentrations, and that VRC, ISC, and PSC may exert similar residual activity against *T. asahii* with longer 24-h exposure.

### 7. Pathogenicity in the *Galleria mellonella* infection model

In the *Galleria mellonella* infection model, the PBS-inoculated group maintained approximately 100% survival throughout the 120-h observation period. Survival curves were compared with the PBS control using the log-rank test. In contrast to our earlier, lower-inoculum experiments, *C. dermatis*, *T. asahii*, *C. albicans*, and *R. oryzae* each showed clear, inoculum-dependent declines in survival at the higher doses used here (Fig. 6). For *C. dermatis* (Fig. 6A), the highest inoculum (1.0×10^6^ CFU/larva) caused a rapid decline in survival, to approximately 72% at 12 h and 22% at 24 h, with survival falling further to approximately 5% by 120 h (*P* < 0.0001 vs. PBS). The intermediate inoculum (5.0×10^5^ CFU/larva) produced a similar, slightly less steep decline, reaching approximately 33% by 24 h and plateauing at approximately 17% by 120 h (*P* < 0.0001 vs. PBS), whereas the lowest inoculum (5.0×10^4^ CFU/larva) produced no mortality and did not differ significantly from PBS. For *T. asahii* (Fig. 6B), the highest inoculum produced a more gradual decline than that seen for *C. dermatis*, reaching approximately 36% survival at 24 h and approximately 9% by 120 h (*P* < 0.0001 vs. PBS). The intermediate inoculum (5.0×10^5^ CFU/larva) showed a comparatively slower decline, reaching approximately 53% by 24 h and plateauing at approximately 29% by 120 h (*P* = 0.0006 vs. PBS), whereas the lowest inoculum (5.0×10^4^ CFU/larva) produced no mortality and did not differ significantly from PBS. For *C. albicans* (Fig. 6C), the highest inoculum (1.0×10^6^ CFU/larva) produced the most rapid decline in survival among the three yeast species tested, falling to approximately 82% at 12 h, 12% at 24 h, and 0% by 96 h (*P* < 0.0001 vs. PBS). The intermediate inoculum (5.0×10^5^ CFU/larva) showed a similar but less steep decline, reaching approximately 12% by 48 h, and 0% by 108 h (*P* < 0.0001 vs. PBS), whereas the lowest inoculum (5.0×10^4^ CFU/larva) produced no mortality and did not differ significantly from PBS. For *R. oryzae* (Fig. 6D), all three inocula produced rapid and severe declines in survival relative to PBS (*P* < 0.0001 for each), with even the lowest inoculum tested (1.0×10^3^ CFU/larva) resulting in complete mortality by 72 h, a substantially smaller inoculum than required to produce comparable mortality with *C. dermatis*, *T. asahii*, or *C. albicans*.

**Fig. 6.**
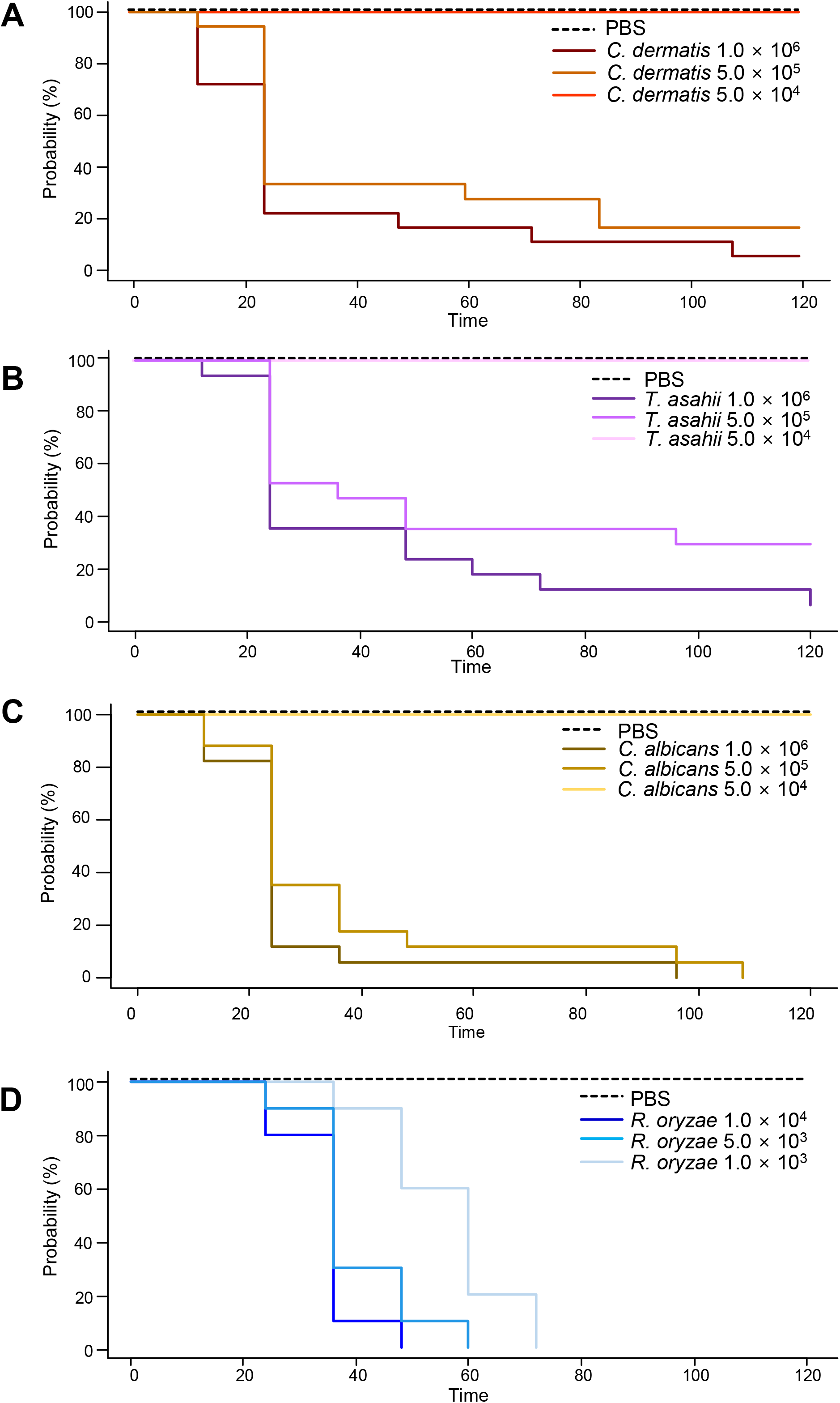
Kaplan–Meier survival curves in the *Galleria mellonella* infection model over 120 **h**. Survival is shown for larvae inoculated with (**A**) *C. dermatis* (5.0×10^4^, 5.0×10^5^, or 1.0×10^6^ CFU/larva), (**B**) *T. asahii* (5.0×10^4^, 5.0×10^5^, or 1.0×10^6^ CFU/larva), (**C**) *C. albicans* (5.0×10^4^, 5.0×10^5^, or 1.0×10^6^ CFU/larva), or (**D**) *R. oryzae* (1.0×10^3^, 5.0×10^3^, or 1.0×10^4^ CFU/larva), or with PBS (control). *Galleria mellonella* larval survival (probability) was monitored every 12 h for up to 120 h. *CFU*, colony-forming unit.

These findings indicate that *C. dermatis*, *T. asahii*, and *C. albicans* each exhibit dose-dependent pathogenicity in the *Galleria mellonella* model when tested at sufficiently high inocula and observed over a longer time course. Among the three yeast species, *C. albicans* produced the most rapid and complete decline in survival at the highest inoculum tested, reaching 0% survival, followed by *C. dermatis*, whereas *T. asahii* showed a somewhat slower and less complete course. Nevertheless, *R. oryzae* was the highest pathogenicity on a per-CFU basis, producing comparable or greater mortality at inocula one to three orders of magnitude lower than those required for *C. dermatis*, *T. asahii*, or *C. albicans*.

## Discussion

In this study, we characterized *C. dermatis* isolated from the sputum and blood cultures of an immunocompromised patient, including molecular identification, pathogenicity, biofilm formation, antifungal susceptibility, drug combination effects, and post-antifungal effects (PAFE). In the *Galleria mellonella* model, *C. dermatis*, *T. asahii*, and *C. albicans* all showed clear, inoculum-dependent pathogenicity at sufficiently high inocula (5.0×10^5^–1.0×10^6^ CFU/larva) over 120 h, although *R. oryzae* remained the most potent pathogen on a per-CFU basis. This is consistent with *C. dermatis* having been recovered from a blood culture and with previously reported cases of *C. dermatis* fungemia in immunocompromised hosts [4, 5].

In line with previous reports indicating that commercial MALDI-TOF MS and biochemical systems frequently fail to differentiate *C. dermatis* from *C. mucoides* [28, 29, 30], the Vitek MS platform could not distinguish our isolate from *C. mucoides*. Indeed, correct identification of this species in earlier clinical cases was achieved only after molecular sequencing [4]. Molecular phylogenetic analysis confirmed the identification independently of the MycoBank similarity search (Table S4), with markedly higher resolution from the IGS1 region (bootstrap 100%) than from ITS alone (63%, decreasing to 43–45% at deeper nodes) [27]. These findings support two-locus sequencing as a more reliable identification approach for this species than phenotype- or MALDI-TOF-based methods alone.

Our initial Galleria experiments, using lower inocula (1.0×10^3^–5×10^4^ CFU/larva) and a shorter (72 h) observation period, showed no decline in survival for *C. dermatis*, *T. asahii*, or *C. albicans* relative to *R. oryzae*. However, increasing the inoculum to 1.0×10^6^ CFU/larva and extending observation to 120 h revealed clear, dose-dependent pathogenicity for all three species, indicating that our earlier findings reflected an inoculum/observation-time threshold rather than a true absence of virulence. Notably, *C. albicans* produced the most rapid and complete mortality at the highest inoculum, followed by *C. dermatis*, whereas *T. asahii* showed a slower and less complete decline, despite *T. asahii* being the more firmly established human pathogen; this graded pattern may in part reflect differences in the *in vitro* growth rates of these species at 37□, particularly the faster growth of *C. albicans* and *C. dermatis* relative to *T. asahii* (Fig. 3E), a temperature close to that used in the Galleria assay. *R. oryzae* nonetheless caused comparable mortality at inocula 1–3 orders of magnitude lower, consistent with its established virulence in mucormycosis. Comparison with the previously reported murine model of *C. dermatis* pathogenicity [10] remains an important direction for future work.

*C. dermatis* is classified as a major antigenic serotype of summer-type hypersensitivity pneumonitis (serotype I) [9], accounting for approximately 41% of *Trichosporon* isolates recovered from the homes of SHP patients [9] and, as with *T. asahii* in serotype II disease [7, 8], its detection in sputum may reflect allergenic potential as well as colonization or pathogenicity. Serological assessment (e.g., anti-*Trichosporon* antibody titers) would help substantiate this in future cases.

*C. dermatis* showed generally strong susceptibility to AmB and the azole agents, intermediate susceptibility to TRB, and intrinsic resistance to MCF, consistent with the known low echinocandin susceptibility of basidiomycetous yeasts [3]. This supports azole monotherapy as a reasonable first-line option.

TRB combined with VRC or ITC showed consistent synergy (FICI ≤ 0.5) across all endpoints, and with PSC at most endpoints. Similar synergy has been reported for *T. asahii* biofilms using sertraline-based combinations [16], but, to our knowledge, this is the first combination-therapy data for *C. dermatis*, supporting TRB and azole combinations as a treatment option for refractory cases.

In contrast to *T. asahii*, where biofilm formation was inhibited by azole agents and AmB but not by TRB or MCF, *C. dermatis* formed a more cohesive biofilm that was more completely inhibited by TRB and AmB. Azole agents achieved only partial inhibition of *C. dermatis* biofilms, with residual biomass ranging from 10% to 20% of control. A modest, non-monotonic increase in *C. dermatis* biofilm biomass at intermediate azole concentrations, most potent for VRC, resembled the paradoxical growth (Eagle) effect reported for echinocandins [37, 38], which has not been widely described for azoles. Established biofilms of both species resisted complete eradication, and eradicating activity did not correlate with inhibitory activity against biofilm formation.

ISC showed the most rapid and pronounced PAFE against *C. dermatis*, evident within 6-h exposure even at low concentrations, whereas the other azoles required 24-h exposure and higher concentrations to achieve comparable suppression of regrowth. This is, to our knowledge, the first report of PAFE data for *C. dermatis*. Although no treatment reports specific to *C. dermatis* are available, ISC has been reported to successfully treat *T. asahii* fungemia refractory to AmB [39], whereas a case of pan-azole-resistant *C. dermatis* causing invasive cholangitis illustrated the therapeutic challenges this genus can pose when standard azole and polyene therapy fails [40]. The present *in vitro* findings, including favorable susceptibility to azole and AmB, a pronounced ISC PAFE, and TRB–azole synergy, may therefore provide a rational basis for treatment selection in future *C. dermatis* infections.

This study has several limitations. First, the isolate was derived from a single case. Second, the *Galleria mellonella* model does not fully reflect the human immune system. Third, allergic sensitization was not directly assessed serologically.

## Conclusion

*Cutaneotrichosporon dermatis*, isolated from the sputum and blood cultures of an immunocompromised patient, was confirmed by ITS/IGS1 sequencing and phylogenetic analysis. *C. dermatis* exhibited dose-dependent pathogenicity in the *Galleria mellonella* model, generally strong susceptibility to azole agents and AmB with intrinsic MCF resistance, and biofilm formation that was more completely inhibited by TRB and AmB than by azole agents, which achieved only partial inhibition even at high concentrations. ISC showed a potent post-antifungal effect, and combination therapy with TRB and an azole agent showed synergy. These findings provide new insight into the biology and clinical treatment of *C. dermatis* infection.

## Acknowledgements

We thank the staff of the Department of Laboratory Medicine at Kumamoto University Hospital for their technical assistance.

## Declarations

### Author contributions

Conceptualization: TY, TN; Methodology: TY, TN; Formal analysis and investigation: TY, TN; Resources: DM; Writing – original draft: TN; Writing – review and editing: DM, JY, YT; Supervision: YT. All authors read and approved the final manuscript.

### Ethics approval

This study was approved by the Ethics Committee of Kumamoto University (Approval No. 2847) as a retrospective analysis of a clinical fungal isolate. The requirement for individual informed consent was waived under an institutional opt-out procedure, as the study analyzed the isolated organism rather than identifiable patient data.

### Consent to participate

This study used a clinical isolate obtained during routine diagnostic testing. The requirement for individual informed consent was waived by the Ethics Committee of Kumamoto University in accordance with an institutional opt-out policy.

### Consent to publish

We confirmed that no identifiable patient information, images, or data are included in this manuscript, consistent with the authors’ decision to omit patient-identifying details.

### Data availability

The ITS and IGS1 sequences generated in this study have been deposited in DDBJ/GenBank under accession numbers [XXXXXXXX] and [XXXXXXXX]. All other data supporting the findings of this study are available from the corresponding author upon reasonable request.

### Declaration of generative AI use

During the preparation of this work, the authors used Claude Sonnet 5 to correct spelling and grammatical errors and to improve English readability. After using this tool/service, the authors reviewed and edited the content as needed and take full responsibility for the content of the published article.

